# Cave-dwelling Planariidae in Croatia exhibit differing levels of cave trait evolution

**DOI:** 10.64898/2026.05.09.723976

**Authors:** Lucija Kauf, Miquel Vila-Farré, Fruszina Ficze-Schmidt, Emilija Bakula, Jochen C. Rink, Helena Bilandžija

## Abstract

The Dinaric karst of Croatia encompasses a network of over 10,000 caves and represents one of the world’s most important subterranean biodiversity hotspots. It is inhabited by remarkably diverse and often endemic species, including planarian flatworms, which are among the rarest macroinvertebrates encountered in cave habitats. Although the presence of cave planarians has long been known, no integrative research on this group has been conducted to date, and the evolutionary relationships between these animals and their surface water counterparts are currently unresolved.

To address these gaps, we combined field sampling, phylogenetic analysis based on *COI* and *18S* genes, and phenotypic characterization. Our results show that cave planariids in Croatia belong to at least three genera and are more widespread and diverse across both Croatia, and the broader Dinaric karst, than previously assumed. We increased the number of cave records in the Dinaric karst from 26 to 37 and documented cf. *Atrioplanaria* and *Phagocata* in Croatian caves for the first time. Phylogenetic reconstructions suggest numerous independent cave colonization events, including multiple instances within the genera *Crenobia* and cf. *Atrioplanaria*. Variation in pigmentation and eye reduction, both within and between populations, further reveal heterogeneous evolutionary trajectories of cave-associated phenotypes.

The biogeographical patterns and high genetic diversity we report here point to a complex evolutionary history of planariids in the Dinarides. Our newly generated molecular phylogenies and systematic documentation of trait variability establish Planariidae as a valuable model for studying mechanisms underlying convergent evolution of pigment loss and eye reduction in cave environments.

## Introduction

Caves and subterranean ecosystems are present around the world, but because access is restricted in comparison to other terrestrial ecosystems, they remain among the least researched habitats. One such understudied region and hotspot for subterranean biodiversity is the Dinaric karst [1], [2], [3], [4], [5], [6], a limestone formation running parallel to the eastern coast of the Adriatic Sea [7]. Due to its high biodiversity both above and below ground, it provides a natural laboratory for investigating evolutionary processes such as speciation, radiation, phenotypic plasticity, and convergence [8], [9], [10], [11], [12], [13], [14]. With respect to the latter process, while cave-dwelling organisms belong to many different phyla, their adaptations to cave environments tend to converge on a shared set of troglomorphic phenotypes [15], the most iconic of which are the loss of body pigmentation and visual senses. These are often accompanied by non-visual sensory enhancements, metabolic rearrangements, and neural and behavioural alterations [15]. Mechanisms leading to convergent evolution of cave-associated traits have been elucidated in only a few examples, with the focus primarily on pigment loss [16], [17], [18], [19].

Given their presence in both subterranean and surface realms of the Dinaric karst [20], [21], [22], [23], [24], [25], [26], [27], [28], [29], [30], although they are amongst the rarest cave macroinvertebrates, and amenability to mechanistic studies of both pigment biosynthesis and eye formation [31], [32], [33], [34], [35], freshwater planarians represent an ideal model for investigating the convergent evolution of cave-related traits. However, this requires resolution of the phylogenetic relationships between surface and cave populations so that appropriate model species can be used for comparative analyses.

Research on planarian species in the Dinaric karst and Croatia dates back to the beginning of the 20^th^ century [20], [21] but with few exceptions (e.g., Knezović et al. [26] and Vila-Farré et al. [36]) has been sporadic and non-systematic [22], [23], [28], [29], [30]. To date, only 16 planarian species have been recorded in Croatia, six of which belong to the family Planariidae [36]. In the Dinaric karst, 26 Planariidae localities originally designated as subterranean have been reported (Appendix 1) [21], [22], [23], [25], [28], [29], [30], [37]. Of these, 22 are caves, and four are springs (two caves and one spring are located in Croatia). Altogether, nine different species and one subspecies, as well as one unidentified Planariid were recorded in these localities: *Atrioplanaria opisthogona* (Kenk, 1936), *Atrioplanaria racovitzai* (Beauchamp, 1928), *Crenobia anophthalma* (Mrazek, 1907)*, Crenobia montenigrina* (Mrazek, 1904), *Phagocata bosniaca* (Stanković, 1926), *Phagocata dalmatica* (Stanković & Komarek, 1927), *Phagocata illyrica* (Komarek, 1919), *Phagocata ochridiana* (Stanković & Komarek, 1927)*, Planaria torva* (Müller, 1773), and *Planaria torva* var. *stygia* (Kenk, 1936) [21], [22], [23], [25], [28], [29], [30], [37]. Only *Ph. dalmatica* and *C. anophthalma* were recorded within the territory of today’s Croatia [22], [37], while one record of *C. montenigrina* from the cave Golubnjača on Ujilica Mountain [28], [29] lacks sufficient information to determine whether it is located in Croatia or Bosnia and Herzegovina (B&H).

Here we report a systematic analysis of biodiversity of cave planariids in Croatia, combining field sampling, phylogenetic reconstructions using mitochondrial *cytochrome oxidase subunit I* (*COI*) and nuclear *18S* genes, and characterisation of hallmark cave traits (namely, reduction or loss of pigmentation and eyes). Resulting molecular phylogenies and patterns of trait variability within and between populations suggest multiple instances of cave colonisation by planariids in the Dinarides and heterogeneous evolutionary trajectories of cave-related traits. Our work reveals greater planarian biodiversity than previously recognised in Dinaric karst ecosystems and sets the stage for future studies using cave Planariidae as a model for understanding mechanisms of convergent evolution.

## Methods

### Sample collection

The cave specimens analysed in this study were collected from 11 localities by our team members and colleagues from the Croatian Biospeleological Society during extensive cave fauna surveys conducted over the past three decades. All the sampling was authorised by the Croatian national authorities (Permit numbers 517-05-1-1-20-05 and 517-10-1-1-22-5). Five surface locations were either specifically targeted for this study or sampled as part of a wider effort to analyze planarian biodiversity [36] (Figure 1A, Figure 1B, Appendix 2). Planarians were sampled using brushes, transfer pipettes, and/or large pipettes [38]. Native water was collected when possible and used to gradually acclimate animals to a standardized formulation of planarian water (PW, Montjuïc recipe) [39] or cave planarian water (cPW, 1:4-diluted PW in RO water). All cave animals were maintained in lab in incubators without light at a constant temperature of 8 or 12 °C and fed monthly with euthanised *Asellus aquaticus* (Linnaeus, 1758). Surface animals were kept in the same conditions but fed with calf liver. Depending on the population, animals were maintained in the lab for a few weeks to a few years.

**Figure 1A.**
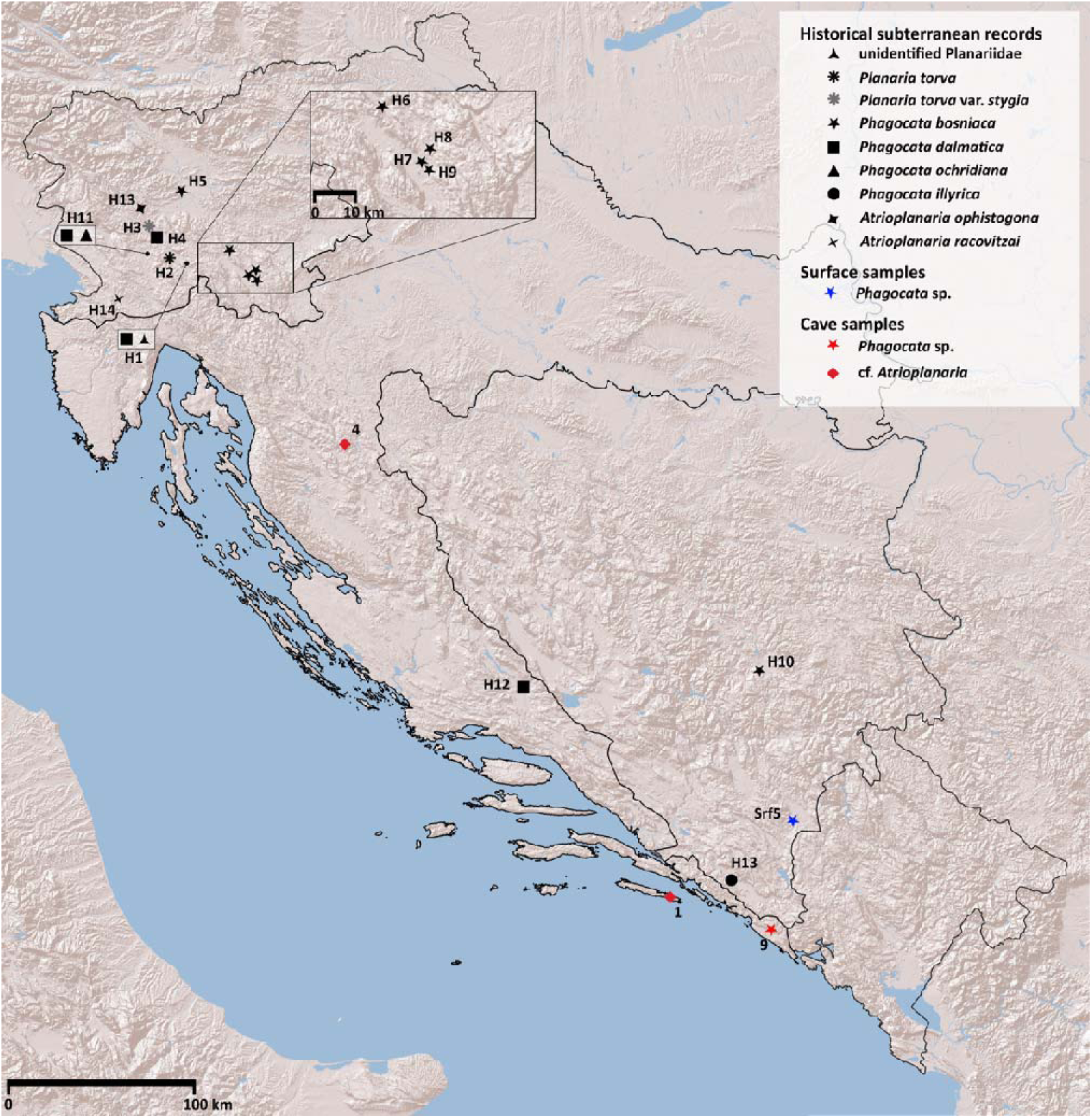
Map of historical and newly sampled localities of *Phagocata* and cf. *Atrioplanaria* in the Dinaric karst.

**Figure 1B.**
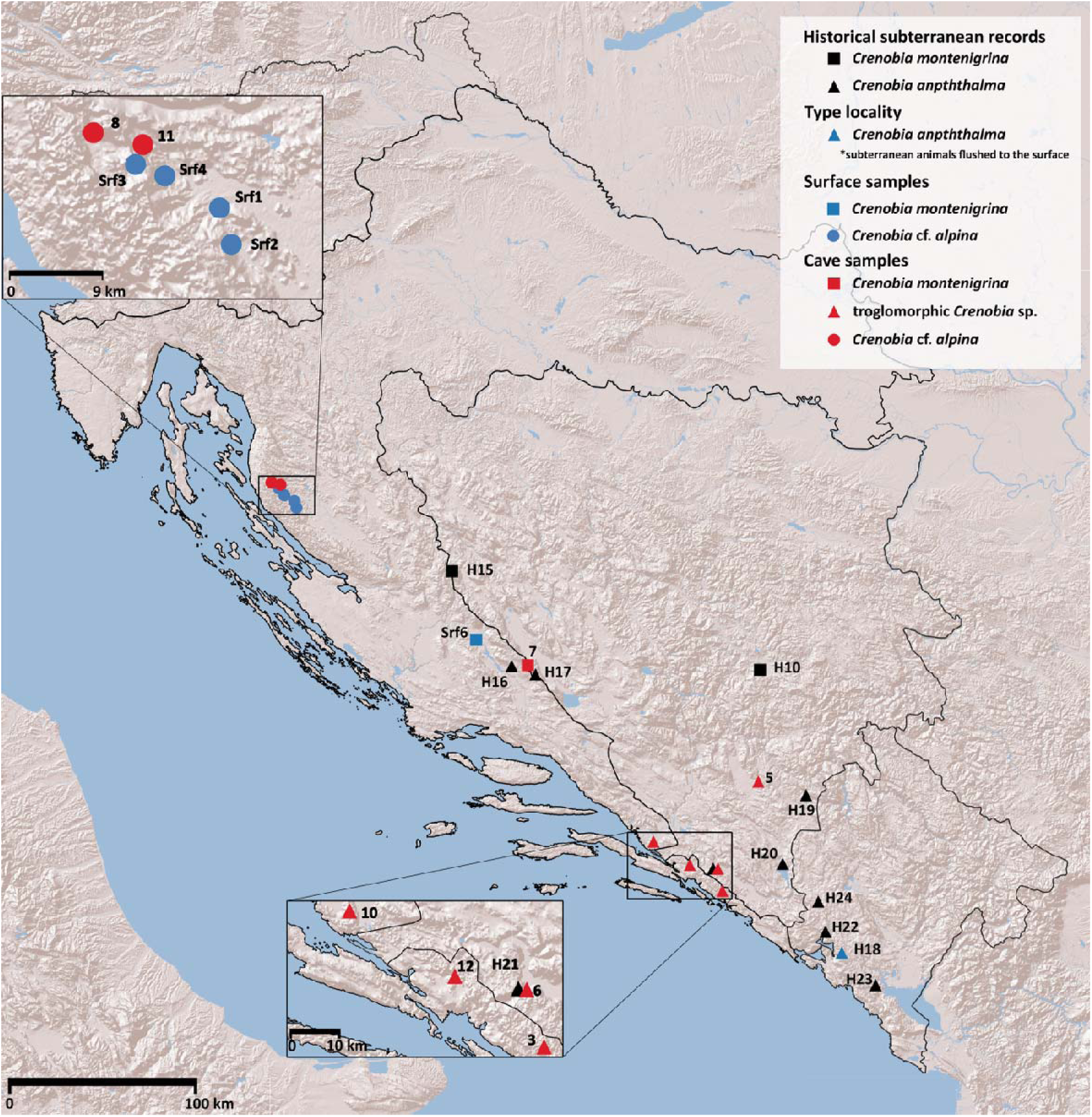
Map of historical and newly sampled localities of *Crenobia* in the Dinaric karst.

### Imaging

Animals were imaged using a Canon EOS 6D Mark II camera equipped with Canon EF 100mm f/2.8L lens, or Canon EDS 250D camera equipped with a Sigma 150mm 1:2.8 APO Macro DG HMS lens for whole habitus images of individuals larger than 2 mm. Images of smaller individuals, as well as detailed images, were taken using either a ZEISS Stereo Microscope Stemi 508 paired with a ZEISS Axiocam 208 colour digital camera or Canon EDS 250D camera mounted on a Motic SZM-171 stereo microscope.

### DNA extraction and PCR

Animals were starved for three weeks or more prior to DNA extraction and were either stored in 96–100% ethanol or processed directly. DNA extraction and *COI* PCR amplification (MVCOI900 primer pairs) either followed published protocols [36] or were performed using commercial DNA extraction kits (Qiagen DNeasy Blood & Tissue Kit, Cat. No. 69504; Zymo Research Quick-DNA Microprep Kit, Cat. No. D3020), followed by a *touch-down* PCR protocol. PCR amplification was conducted using commercial Taq MasterMixes (highQu Cat. No. HSM0301; Promega Cat. No. M7832). *Touch-down* PCR was performed with an initial denaturation of 5 min at 95°C, followed by seven cycles of 30 s at 95°C, 30 s at 60°C (decreasing by 2°C over the first seven cycles), and 1 min at 70°C, followed by 30 cycles with annealing at 46°C. Final extension was performed for 3 min at 70°C.

*18S* sequences were obtained by PCR amplification using NZYTaq II 2× Green Master Mix (Cat. No. MB35803) with 18S_F1 and 18S_R9 primers [40], [41]. The reaction conditions were: (1) 5 min at 95°C, (2) 30 s at 95°C, (3) 30 s at 45°C, (4) 2 min at 68°C, and (5) 5 min at 68°C. Steps 2–4 were repeated for 30 cycles. All commercial kits were used according to the manufacturers’ instructions.

Amplification products of both genes were analysed by gel electrophoresis on a 1 % agarose gel and purified using the QIAquick PCR Purification Kit (Cat. No. 28104). The purified PCR products were Sanger sequenced using the amplification primers in both directions, with additional 18S_4F and 18S_5R [40] for 18S by Macrogen. Low-quality bases were trimmed and sequences were assembled in Geneious Prime (version 2026.0.2) [42]. Predicted protein sequences were analyzed to ensure no internal stop codons were present, and coding sequences were then used in BLASTN [43] queries against the NCBI database [44].

### Extraction of sequences from transcriptomes

Our dataset was complemented with additional Planariidae sequences from Planmine [45], [46], as well as from unpublished transcriptomes of cave planarians generated by our group (Appendix 2) via a BLAST search integrated in Geneious Prime [42]. The hits were manually inspected, and regions homologous to the PCR-obtained *COI* and *18S* fragments were extracted. In cases where both DNA-amplified and transcriptome-retrieved sequences were available for the same population, the transcriptome sequences were used.

### *COI* and *18S* NCBI search

In order to investigate the relationships between the samples and of those with published sequences, the NCBI nucleotide database [44] was searched for both genes on 15.01.2026. The search was limited to the genera *Crenobia*, *Phagocata* and cf. *Atrioplanaria* based on the preliminary NCBI BLAST [43] results and the transcriptome-based phylogenetic trees in [36], [38]. Additionally, sequences from the planariid *Polycelis felina* (Dalyell, 1814) were used as an outgroup because of its phylogenetic position [36], [38]. Only the samples with both *COI* and *18S* were retained for subsequent analyses.

### Sequence alignment and phylogenetic tree reconstruction

In total, 72 sequences were analysed (16 generated in this study, seven extracted from published transcriptomes, and 49 retrieved from NCBI). Sequences of individual genes were aligned using MAFFT (v7.490) [47] (Algorithm G-INS-I; Scoring matrix 200PAM/k=2; Gap open penalty = 1.53; Offset values = 0.123) integrated in Geneious Prime [42]. For the *COI* gene, the alignment was 1,047 bp long, containing 39.20% missing data, 0.22% ambiguities, and 0.04% gaps. The *18S* alignment was 1,823 bp long, containing 46.66% missing data, 0.03% ambiguities, and 1.06% gaps. The missing information in the alignment ends was substituted with Ns, and the sequences were concatenated into a single alignment using the integrated concatenation option in Geneious Prime [42], generating a new alignment with a length of 2,870 bp, and a total of 43.98% of missing data, 0.10% of ambiguities, and 0.69% of gaps, further referred to as ‘full’ alignment. To assess the effect of missing data on phylogenetic relationships, we constructed two additional trimmed alignments using an approach similar to that of Vila-Farré et al. [36]. Positions in the alignment ‘gap50’ have an occupancy of 50% or more, with a length of 1,092 bp and 5.39% of missing data, 0.19% of ambiguities, and 0.03% gaps. The ‘trimmed’ alignment was obtained manually by trimming the alignment ends to include a region in which all sequences contained information. The final length of the ‘trimmed’ alignment was 782 bps, with 2.19% gaps and 0.16% ambiguities (no missing data). The reading frame for the *COI* gene was preserved in both alignments. Maximum Likelihood (ML) phylogenies [48] were inferred from all three alignments using IQ-TREE (version 3.0.1) [49]. Alignment partitions were specified for *COI* codon positions 1, 2, and 3, and for the *18S* gene. Model selection was restricted to models supported by MrBayes, followed by tree construction using the parameters ‘-m MFP -mset mrbayes’. Branch support was assessed with 10,000 ultrafast bootstrap replicates (‘-bnni -bb 10,000’). The branches with ultra-fast bootstrap values under 95 were subsequently collapsed in iTOL (version 7.5) [50], using the ‘delete branches’ function. The same pipeline was used for phylogenetic analyses of each gene individually, with an appropriately adjusted partition file for *COI*, and exclusion of the partition file for the *18S* gene.

Based on ML results, we performed Bayesian inference (BI) on ‘full’ alignment only, since both ‘gap50’ and ‘trimmed’ showed a lack of resolution or relationships inconsistent with transcriptome-based phylogenetic relationships from Vila-Farré et al. [36], [38] (data not shown). Single gene trees were also run for the full length of *COI* and *18S* independently. As determined by IQ-TREE [49] the best models compatible with MrBayes for *COI*+*18S* alignment were: GTR+F+G4 for *COI* codon position 1, HKY+F+G4 for *COI* codon position 2 and GTR+F+I+G4 for *COI* codon position 3, and HKY+F+I+G4 for *18S* partition. MrBayes (version 3.2.7) [51], [52] was run in two independent runs with four chains for each of the alignments. Number of generations was 10,000,000, with a burn-in of 10,000, sampled every 1,000 generations. Chain convergence was assessed first by observing the average standard deviation of split frequencies (<0,01 for all alignments), as well as inspecting trace plots in Tracer (version v1.7.2) [53]. The effective sample size (ESS) for all model parameters exceeded 625, as confirmed using the convenience package (version 1.0.0) in R (version 4.4.3) [54]. Trees were created using the ‘-allcompat’ function in MrBayes [51], [52], and all branches with posterior probability lower than 0.95 were deleted as in ML trees.

## Results

### Preliminary sequence identification

Based on external morphology, some collected populations were preliminarily assigned to the genus *Crenobia*, while the remaining specimens were left unidentified. BLASTN searches [43] of *COI* sequences from putative *Crenobia* showed percentage identity (ID%) values ranging from 86.29% (cave12) to 93.40% (Srf2), with query coverage ranging from 30% (Srf2) to 66% (cave12) relative to *Crenobia* sequences in NCBI. Hits for the remaining samples showed lower ID% values. All *18S* sequences returned *Phagocata* as the best BLAST match, with ID% values ranging from 93.59% to 97.46, including specimens that showed high similarity to *Crenobia* in the *COI* search. Although NCBI contains *Crenobia 18S* sequences [55], these are substantially shorter than those generated in this study, resulting in query coverage below 30%. This size difference, together with the relatively slow evolutionary rate of *18S* [40], [56], [57], likely explains the inconsistent BLAST assignments.

### Phylogenetic relationships

To further characterise cave planariid diversity, we conducted BI and ML analyses of individual *COI* and *18S* datasets, as well as concatenated *COI+18S* alignments from specimens collected in 11 cave and five surface localities. Analyses based on individual genes, trimmed datasets, and all ML reconstructions either lacked resolution or recovered relationships inconsistent with previously published transcriptome-based phylogenies [36], [38] (Appendix 3–7). Therefore, subsequent interpretations were based on the BI analysis of the concatenated *COI+18S* dataset (Figure 2).

**Figure 2.**
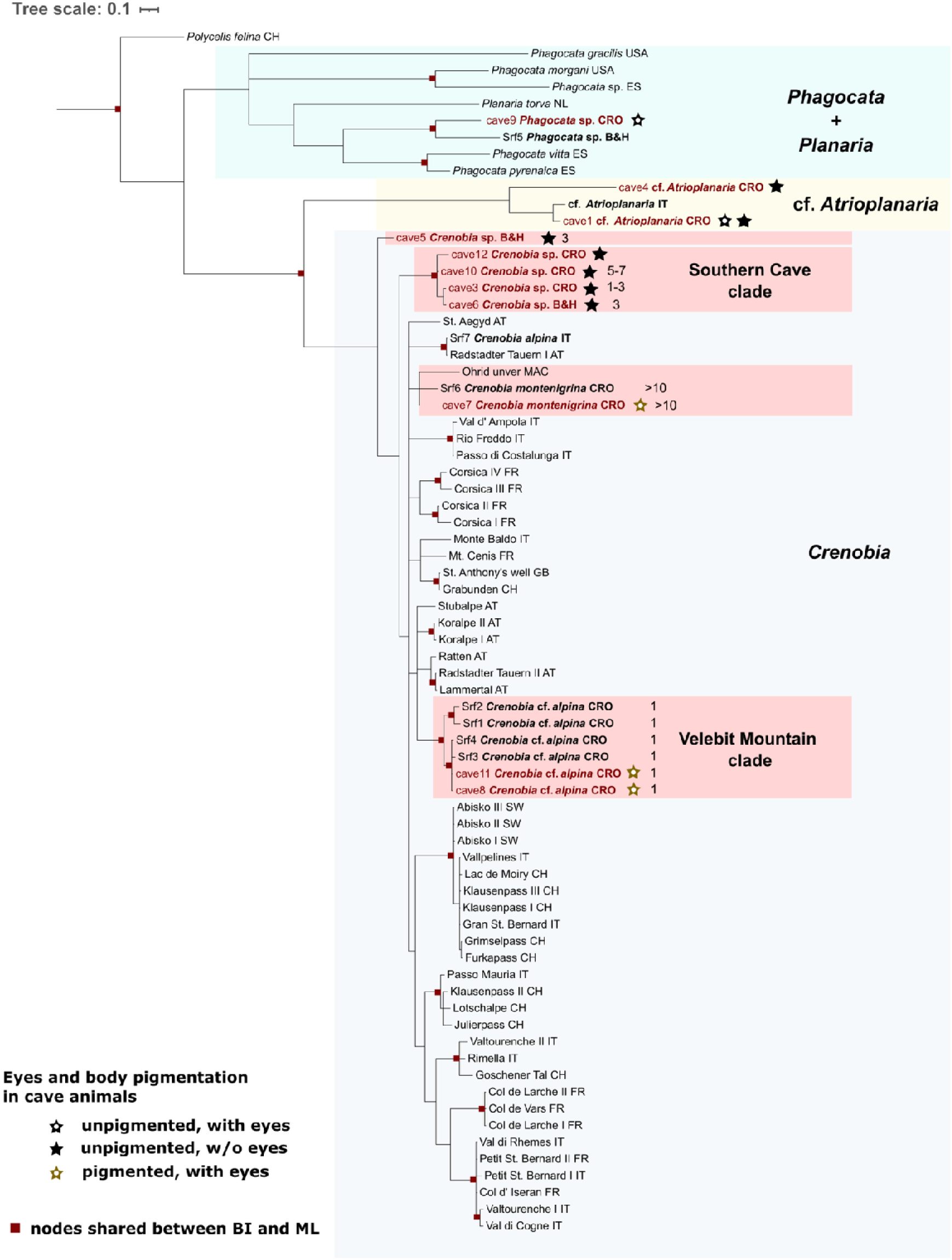
Bayesian inference phylogenetic tree based on the ‘full’ alignment. Branches with posterior probability lower than 95 deleted. Terminals corresponding to cave animals are shown in red. Where available, the number of pharynges is indicated to the right of species names. Colour shadings indicate the retrieved groups. Red shading indicates clades within *Crenobia* that include cave populations.

This analysis identified cave specimens belonging to three genera: *Phagocata*, cf. *Atrioplanaria*, and *Crenobia*. Specimens from cave9 and Srf5 clustered with European *Phagocata*, and formed a sister group to *P. torva*, consistent with previous transcriptomic studies. Sample *Phagocata* sp. ES grouped outside of European lineages and clustered within American taxa [36], [38] (Figure 2).

Specimens from cave1 and cave4 clustered with a previously described surface population from Sardinia [36], [38], and were therefore assigned to cf. *Atrioplanaria*. Their sister clade was *Crenobia*, which included specimens from eight caves and four surface localities. Relationships within *Crenobia* remained only partially resolved due to low statistical support for several nodes. Nevertheless, cave specimens formed four distinct lineages: (1) cave5, which occupied a sister position to all remaining *Crenobia*; (2) the Southern Cave clade, comprising specimens from cave3, cave6, cave10, and cave12; (3) cave7, which clustered with *Crenobia montenigrina*; and (4) cave8 and cave11, which grouped with nearby surface populations from Velebit Mountain (Figure 2).

### Pigmentation and eye variation

As lack of body pigmentation and eyes are the most prominent characteristics of cave-adapted animals, we examined their variation in our animals. Both cave and surface *Phagocata* were unpigmented but possessed well-developed eyes, consistent with previous reports of unpigmented epigean *Phagocata* [58], [59] (Figure 3). The two cf. *Atrioplanaria* populations were also unpigmented. Animals from cave4 were eyeless, while individuals from cave1 showed variability in eye number, ranging from none to two eyes (Appendix 8). Within cave *Crenobia*, the two early-branching groups (cave5 and the Southern Cave clade) lacked both body pigmentation and discernible eyes (Figure 3). In contrast, *C. montenigrina* from cave7 and both cave populations from the Velebit Mountain clade retained well-developed eyes, but showed varying levels of body pigmentation (Figure 4; data not shown).

**Figure 3.**
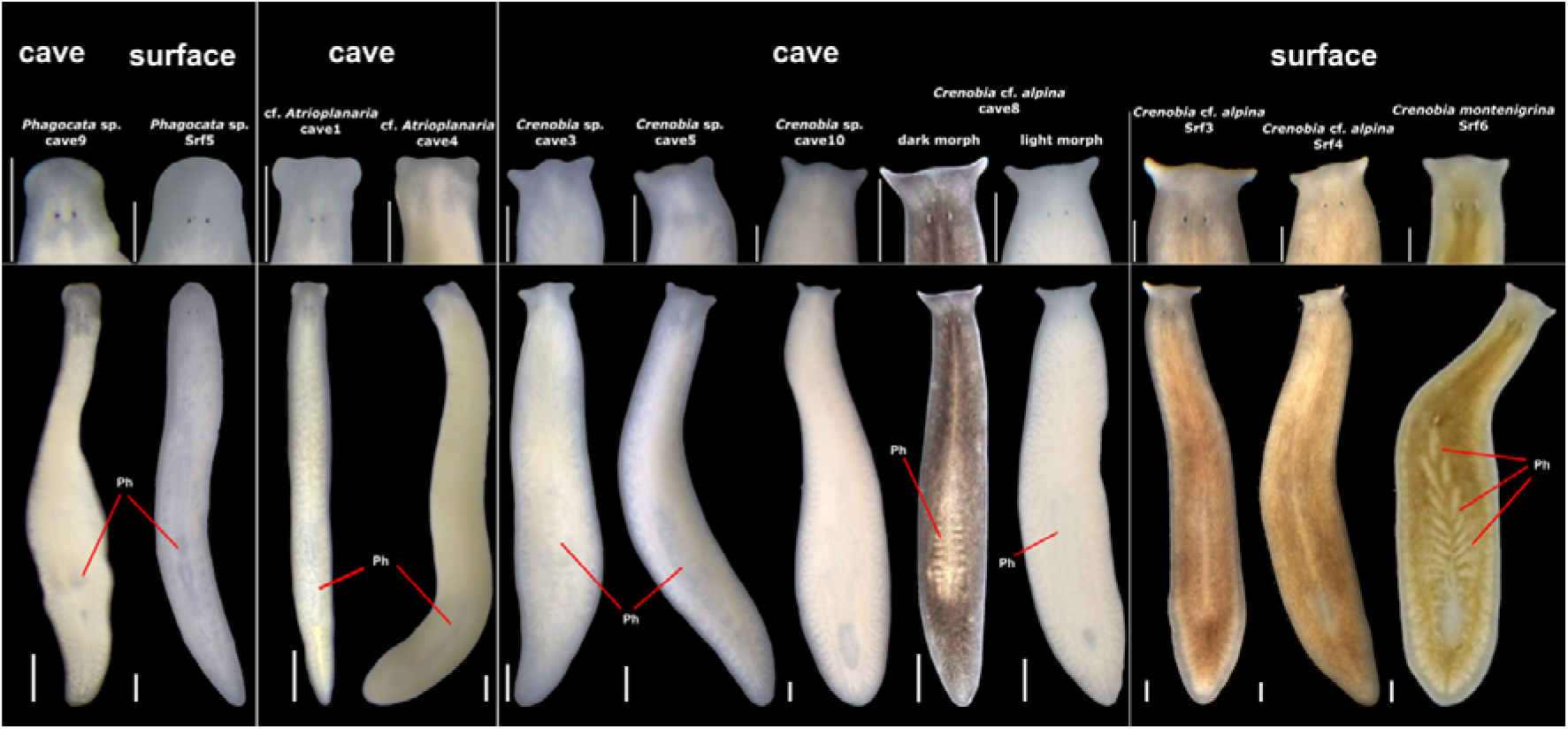
Live specimens of planariids collected during the expeditions. Top panel, anterior region (head) in close-up; bottom panel, whole-body habitus. Ph: pharynx/pharynges. Scale bar, 500 µm.

**Figure 4.**
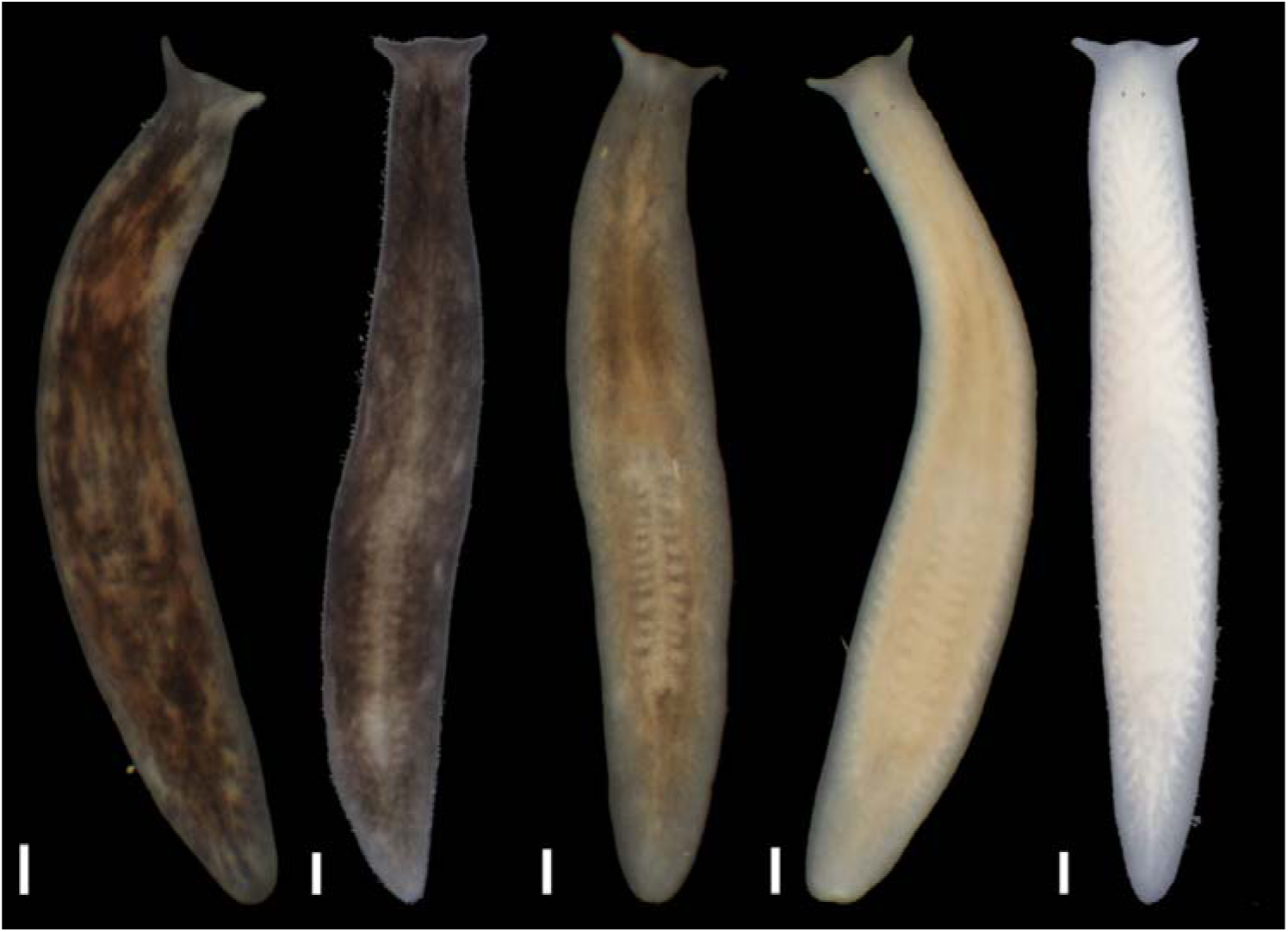
Intrapopulation variation in body pigmentation among living cave8 animals imaged under identical illumination conditions. Scale bar, 500 µm.

### Variability of pharynges number in *Crenobia*

To further characterise *Crenobia* diversity, we examined the number of pharynges, given the established taxonomic value of this trait [37]. All animals from the Velebit Mountain clade were monopharyngeal (both cave and surface individuals), consistent with the description of *C. alpina,* but also *C. bathycola* (Steinmann, 1911) [60] and *C. corsica* (Arndt, 1922) [61]. The remaining specimens were polypharyngeal. Animals from cave7 possessed more than 10 pharynges, consistent with previous descriptions of *C. montenigrina* [20], [37], [62], [63]. Animals from cave5 and the Southern Cave clade externally resembled *C. anophtalma*, the only known obligatory cave *Crenobia* species [21], [37], in their complete absence of pigmentation and eyes. However, they showed variability in the number of pharynges. While specimens from cave5 (N=4), cave3 (N=3) and cave6 (N=1) possessed up to three, animals from cave10 (N=5) had between five and seven pharynges (Figure 2, 5). Animals from cave12 were not examined due to the lack of material.

**Figure 5.**
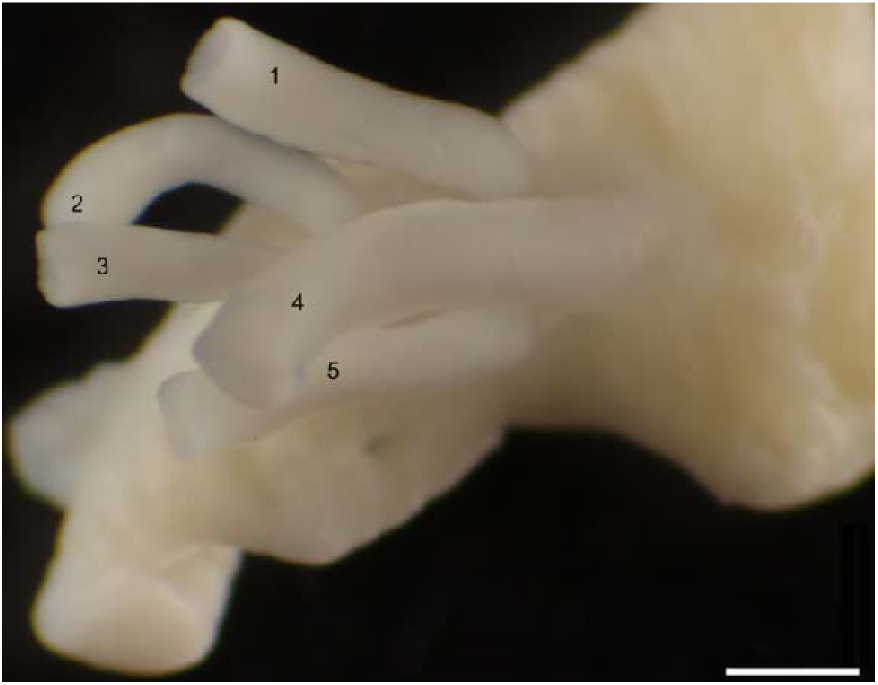
Expelled pharynges from an individual collected in cave10 after fixation in 96% ethanol. Most animals had five pharynges, with some having six or seven.

## Discussion

Our sampling efforts revealed at least seven different Planariidae lineages in the caves of the Dinaric karst: two unidentified cf. *Atrioplanaria*, one unidentified *Phagocata*, and four *Crenobia* (Figure 2). We provide the first cave record of *Phagocata* in Croatia, as the previously reported subterranean record of *Ph. dalmatica* in Croatia was collected in a spring [22]. Similarly, *C.* cf. *alpina* and *C. montenigrina* are documented here from Croatian caves for the first time, and cf. *Atrioplanaria* for the first time in the country. This substantially increases the known subterranean diversity of planariids, from a single species (*C. anophtalma*) and two cave localities to three genera across 11 caves. Given that planarians are rarely encountered during cave sampling in the Dinaric karst, this increase is particularly notable and underscores the species richness of planarian fauna in Croatia [36], as well as the Dinaric karst.

Our phylogenetic analysis on the combination of the rapidly evolving mitochondrial *COI* gene and the more slowly evolving nuclear *18S* gene [40], [56], [57] was sufficient to produce an overall tree topology concordant with the trees based on transcriptomic data [36], [38]. The results obtained clarify the evolutionary relationships of all three detected genera and reveal distinct biogeographical patterns. The placement of Srf5 and cave9 within European *Phagocata* confirms cave9 as the first true cave record of this genus in Croatia. The grouping of the European *Phagocata* lineages with *P. torva,* and the separation from American *Phagocata* lineages, with the exception of the likely misassigned *Phagocata* sp. ES, is expected, as these relationships were also recovered in previously published analyses [36], [38]. The intriguing phylogenetic proximity between cf. *Atrioplanaria* from the island of Mljet (cave1; one of the southernmost Croatian islands) and a Sardinia population [38], rather than to specimens from the geographically closer locality in Croatia (cave4 population), is statistically supported in all Bayesian analyses and ML using *18S* (Figure 2, Appendix 4, 5, 7, 9). Given the substantial distance between the two caves and their association with distinct Croatian river catchment systems (the Adriatic and the Danube, respectively), a single cave colonisation event by cf. *Atrioplanaria* followed by subterranean dispersal is unlikely. A more plausible scenario is that the Sardinian and Mljet populations are descendants of a lineage that was widely distributed across the Mediterranean during the Messinian salinity crisis [64], [65], while the population from cave4 might have arisen from an earlier split. Comparable trans-Adriatic paleogeographic patterns have been reported in *Niphargus* amphipods [66] reflecting the role of paleogeographic processes in shaping current distributions of cave fauna.

Variation in pigmentation and eye reduction reveal complex patterns of cave-trait evolution. In *Phagocata* and cf. *Atrioplanaria,* the absence of pigmentation cannot be interpreted as a result of cave adaptation, as unpigmented populations are also known from surface environments [38], [58],[59]. Variation in eye number between and within cf. *Atrioplanaria* cave populations (Figure 3; Appendix 8) suggest either ongoing or heterogeneous evolutionary processes. Similar to the case in cave1, eye-phenotype variability has recently been documented in *Girardia multifiverticulata* de Souza, Morais, Cordeiro & Leal-Zanchet, 2015 [34], [67]. Such variability could reflect differences in the stage of adaptation to cave environments, since the loss of eyes has been associated with the time elapsed since cave colonisation [68] and eyeless cave *Atrioplanaria* have previously been documented, e.g., in Slovenia [23]. However, such interpretations should be treated cautiously, as the relationship between trait loss and time since colonization is not necessarily linear [9], [69], [70], [71], [72].

The existence of four cave *Crenobia* lineages strongly suggests multiple independent cave colonisation events within the genus. Although relationships within *Crenobia* remain only partially resolved, we recovered all well-supported relationships presented by Brändle et al. [55]. The cave5 and Southern Cave clades are formed by cave-adapted individuals with a distant phylogenetic position relative to the surface population (Figure 2, Appendix 7, 9), which is consistent with the climatic relic hypothesis of cave colonisation [73], [74], [75], [76]. As such, they represent remnants of formerly widespread surface populations that escaped extinction during past adverse climatic periods (e.g., glacial periods) by finding refuge in caves. In contrast, the Velebit Mountain and *C. montenigrina* clades contain both cave and surface populations, which is more consistent with the adaptive shift hypothesis of cave colonisation. Under this hypothesis, cave colonisation is primarily driven by ecological opportunity, where surface populations colonise the empty niche (in this case, the cave environments), resulting in closely related surface and cave populations or species [15], [77], [78], [79].

Trait variability in pigmentation and eye number is remarkable in *Crenobia*. Fully troglomorphic lineages (cave5 and Southern Cave clade) represent the most advanced degree of trait reduction in this genus. Together with the lack of surface relatives, this may indicate the oldest cave colonisation event among the analysed *Crenobia*. In contrast, partial reduction of body pigmentation (pale colour with a pink hue) in cave7 (*C. montenigrina*) in relation to the nearby surface population (data not shown) may indicate a more recent colonisation. Interestingly, the historical record of *C. montenigrina* from the cave Golubnjača references animals that are ‘snow-white, with degenerate eyes’ [29]. These differences in trait evolution within a single species may reflect independent cave colonisation events or divergent selection pressures in different caves, an issue that might be resolved with further species sampling. Finally, the most extensive intrapopulation variability in pigmentation among cave *Crenobia* was recorded at Velebit Mountain (two populations; cave8 shown in Figure 4). Since functions of body pigmentation, such as protection against solar radiation, mimicry, or intraspecific communication, are obsolete in the absence of light, the observed high pigmentation variability might be a result of relaxed selective pressures [80], and reflect a recent cave colonisation. Alternatively, this variability could reflect ongoing gene flow from surface populations [15]. However, since no planarians were found in a nearby surface stream leading to Ponor Štirovača (cave8), despite repeated sampling efforts, this explanation is unlikely.

Beyond evolutionary patterns, our results raise taxonomic questions within *Crenobia*. Sluys [37] recently confirmed the species status of *C. alpina, C. corsica, C. batycola, C. montenigrina,* and *C. anopththalma.* To date, only *C. anophthalma* has been reported with certainty from Croatian subterranean environments [37]. However, our phylogenetic analyses indicate substantially greater diversity. Specifically, we present the first subterranean record of *Crenobia montenigrina* for Croatia, and the third record in the Dinaric karst [22], [29]. Similarly, individuals from the Velebit Mountain clade (four surface and two cave populations) are related to a mostly epigean European species, *C. alpina*. To our knowledge, these specimens represent the first record of a monopharyngeal cave *Crenobia* in the Dinaric karst. Finally, specimens from cave5 and most populations in the Southern Cave clade (cave3, cave6 and cave12) are morphologically similar to *C. anophthalma* [21], [37], [81], with the exception of cave10 individuals having five to seven pharynges (Southern Cave clade). The broad morphological concordance, the occurrence of the two clades within or close to the known range of the species [28], [29], [37], and the existence of a historical *C. anophthalma* record at less than 600 m from cave6 [28], [29] all support the assignment of those lineages to *C. anophthalma.* If cave10 individuals belong to *C. anophthalma*, the number of pharynges may be taxonomically less informative than previously assumed. The phylogenetic position of cave5 specimens, separated from the Southern Cave clade animals, and the higher number of pharynges in cave10 individuals support the existence of additional subterranean species of *Crenobia* in the area. In either case, our findings highlight the need for an integrative taxonomic re-evaluation of *Crenobia* [36].

## Conclusions

Our combined sampling, phylogenetic, and trait-based analyses demonstrate that cave planariids are more widespread and diverse in Croatia as well as the Dinaric karst than previously recognised. The emerging biogeographical patterns, shaped by both historical and ecological processes, point to a complex evolutionary history and underscore the need for further research in this area. The newly generated gene sequences, especially for *Crenobia*, provide a foundation for future barcoding and integrative taxonomic studies. The occurrence of multiple cave colonization events, even within the same genus, and availability of multiple lineages exhibiting varying degrees of trait evolution position Planariidae as an ideal system for investigating mechanisms underlying convergent evolution in cave environments.

## Data Accessibility Statement

All newly reported gene sequences are in the process of submission to ENA public repository. Bioproject accession PREJB112134.

## Competing interests

The authors declare that they have no competing interests.

## Funding

This project was funded by the Max Planck Society (J.C.R.) and by the Tenure Track Pilot Programme of the Croatian Science Foundation, Ecole Polytechnique Fédérale de Lausanne, and Project TTP-2018-07-9675 Evolution in the Dark, with funds from the Croatian-Swiss Research Programme (H.B.).

## Authors’ contributions

Wet lab: LK, MVF, FFS, EB. Analyses: LK. First draft: LK with inputs from HB and MVF; Sampling: LK, MVF, JCR, HB. Funding: HB, JCR. All authors read and approved the final version of the manuscript.

## Acknowledgements

This study would not have been possible without the extensive biospeleological sampling efforts conducted by the Croatian Biospeleological Society (CBSS) over the past three decades. We thank Teo Delić, Marko Lukić, Magdalena Grgić, Lada Jovović, Jana Bedek, Martina Pavlek, Branko Jalžić, Nikolina Kuharić, Alen Kirin, Hrvoje Cvitanović, Nataša Cvitanović, Tin Rožman, Igor Božić, Filip Šarc, Nina Trinajstić, and all other CBSS members for their fieldwork support. We also thank Kerstin Raab for her support in collecting references. We thank the animal care takers of the MPI-NAT animal services team and Jason Pellettieri for proofreading the manuscript.

## Appendixes and legends

## Appendix 1. Historical records of the subterranean Planariidae in the Dinaric karst. This appendix has also been uploaded separately in the submission system

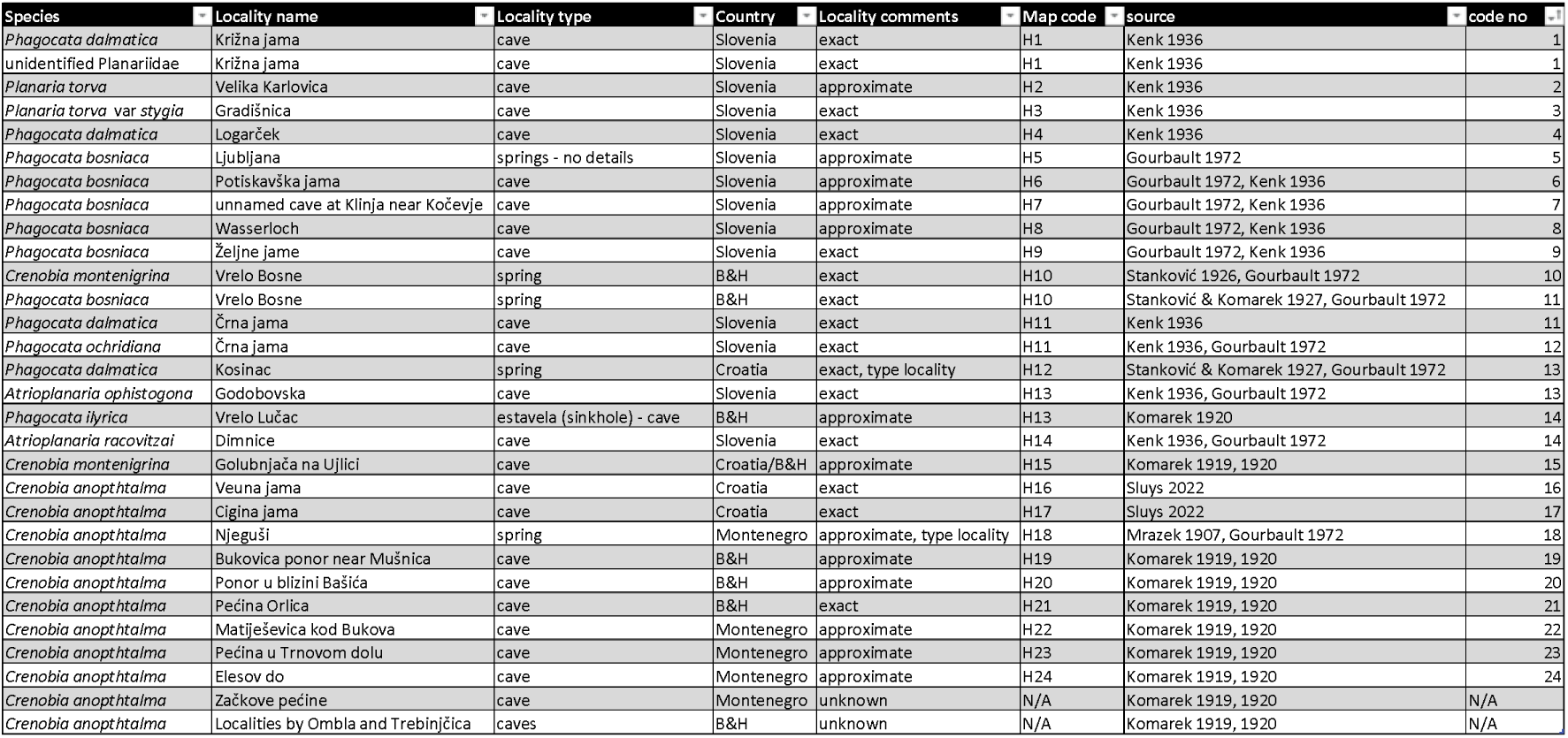

## Appendix 2. List of localities and samples used in the phylogenetic analyses. This appendix has also been uploaded separately in the submission system

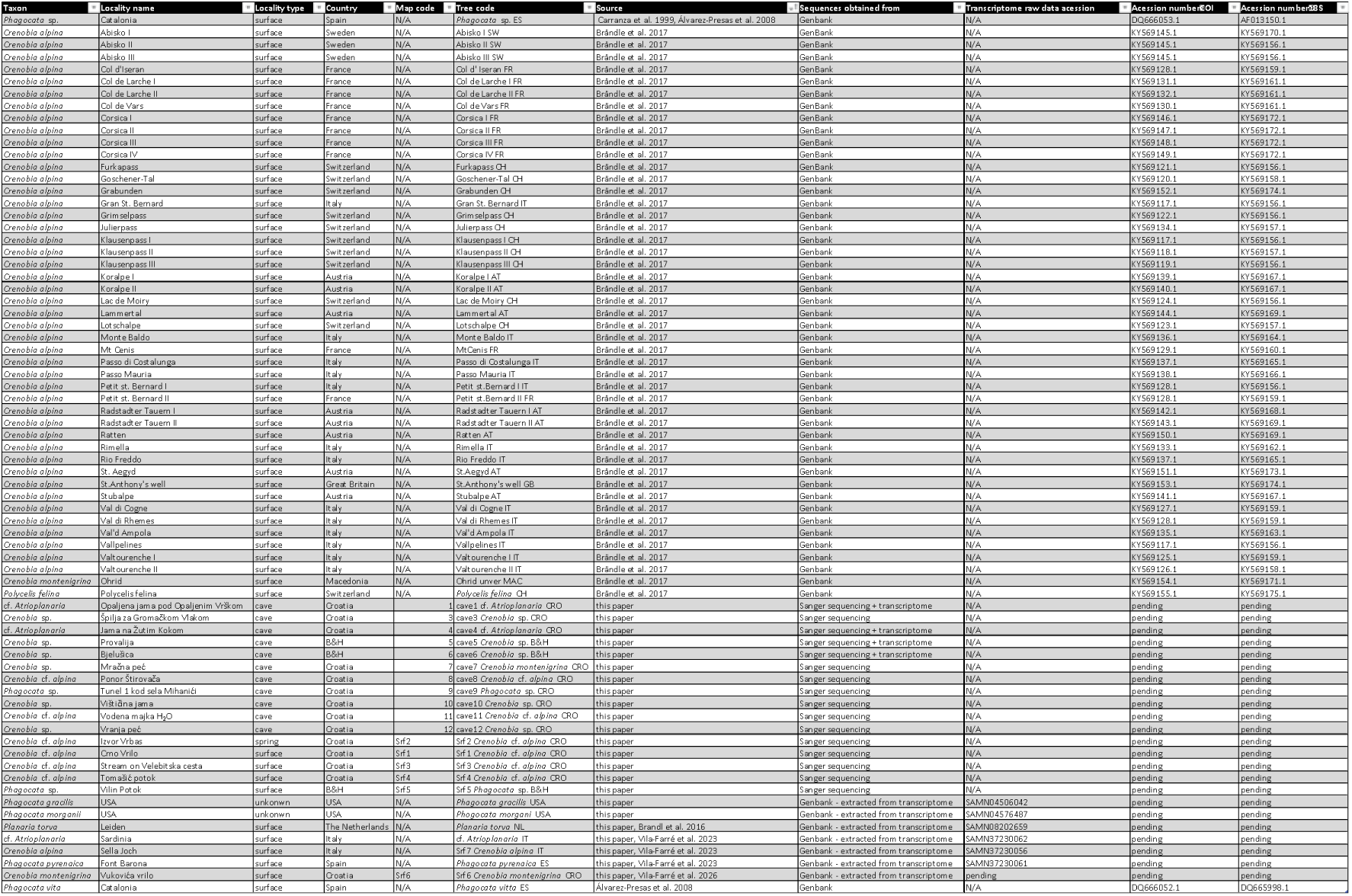

## Appendix 3. ML phylogenetic tree inferred by IQ-TREE from concatenated *COI + 18S* genes, using ‘full’ alignment dataset. Bootstrap values lower than 0.95 were collapsed using the ‘delete’ function in iTOL

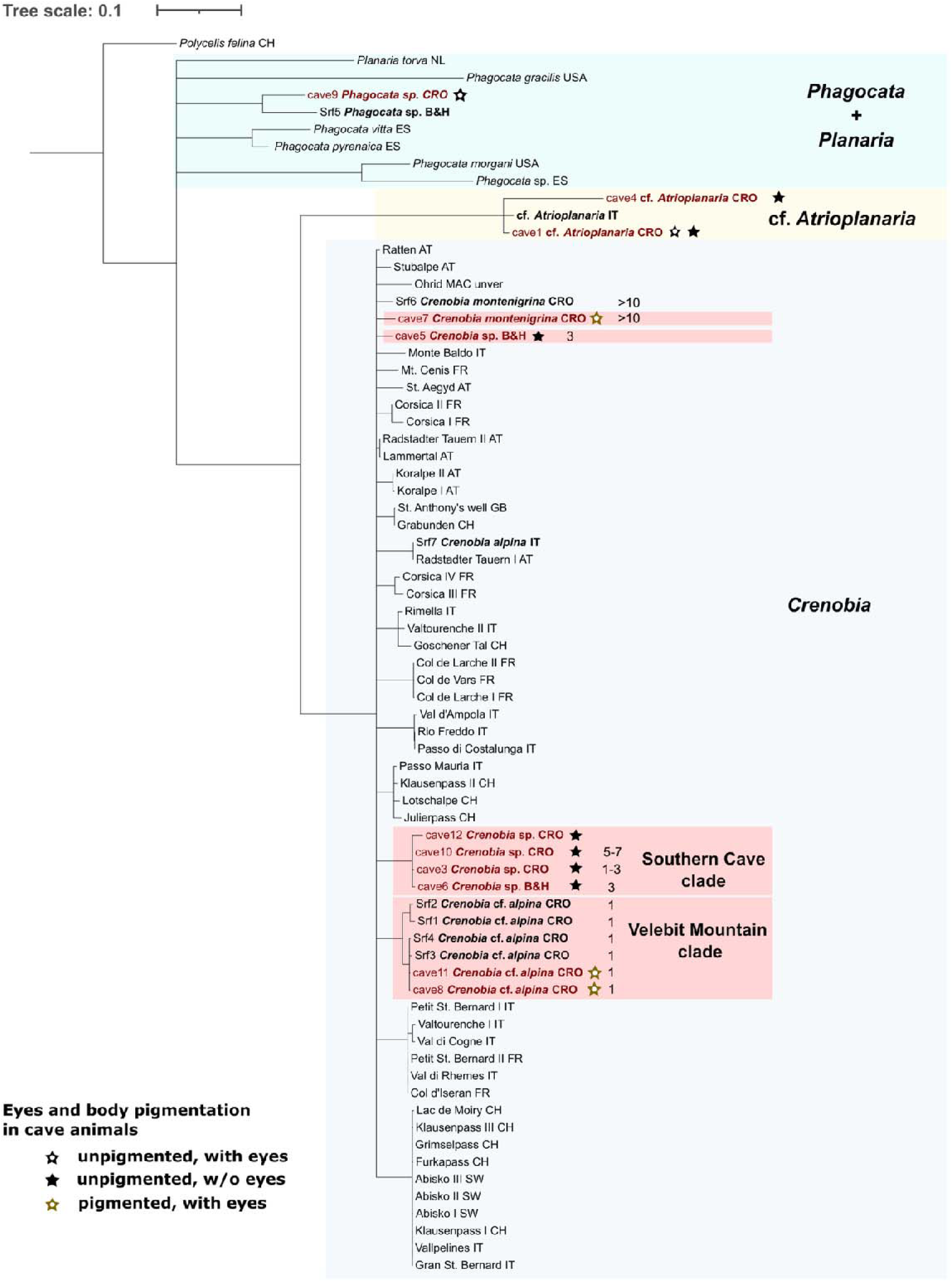

## Appendix 4. ML phylogenetic tree inferred by IQ-TREE from *18S* gene only, using the 1,823 bp long sequence alignment. Bootstrap values lower than 0.95 were collapsed using the ‘delete’ function in iTOL

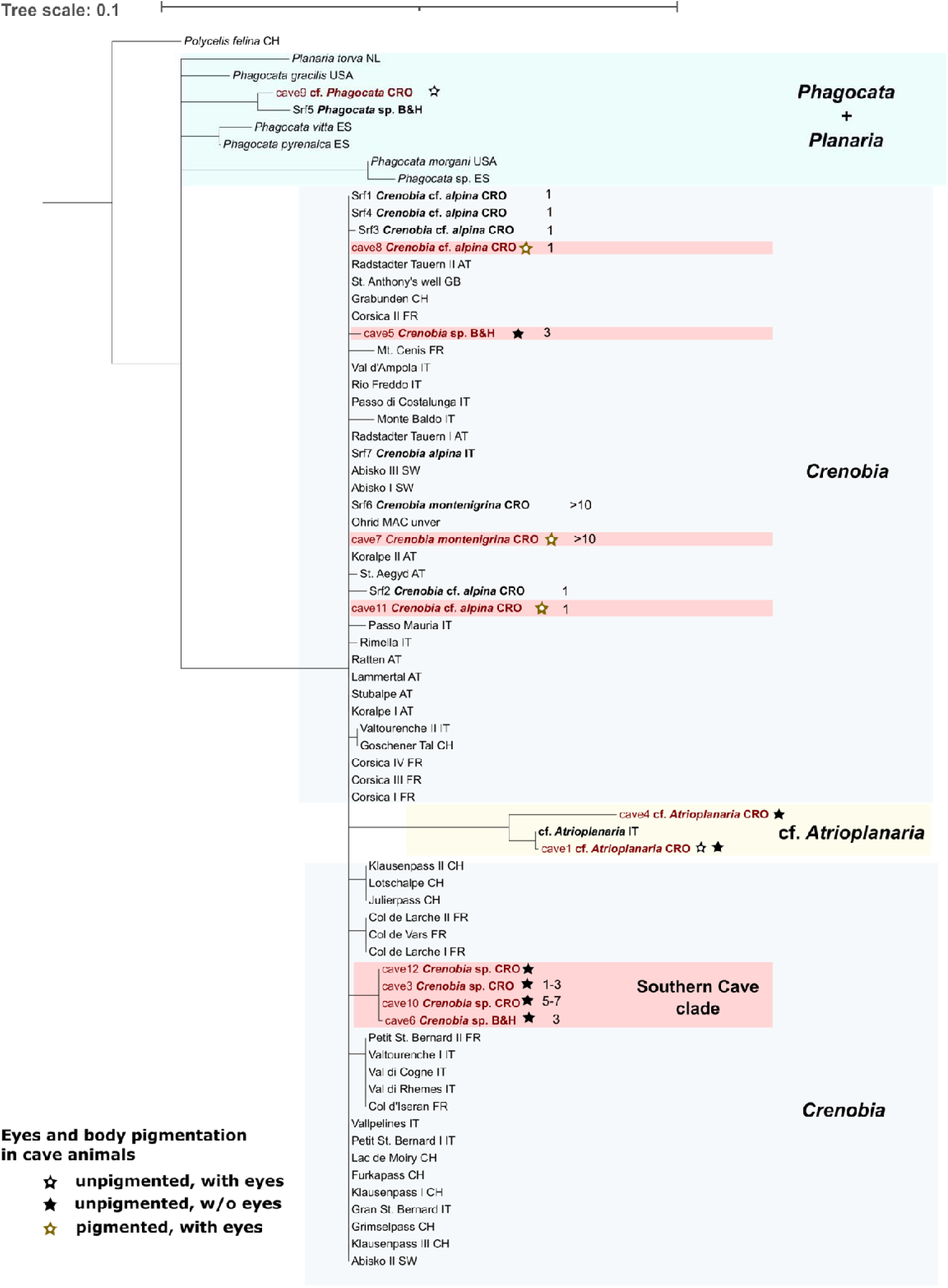

## Appendix 5. BI phylogenetic tree inferred by MrBayes from *18S* gene, using the 1,823 bp long sequence alignment. Posterior probability values lower than 0.95 were collapsed using the ‘delete’ function in iTOL

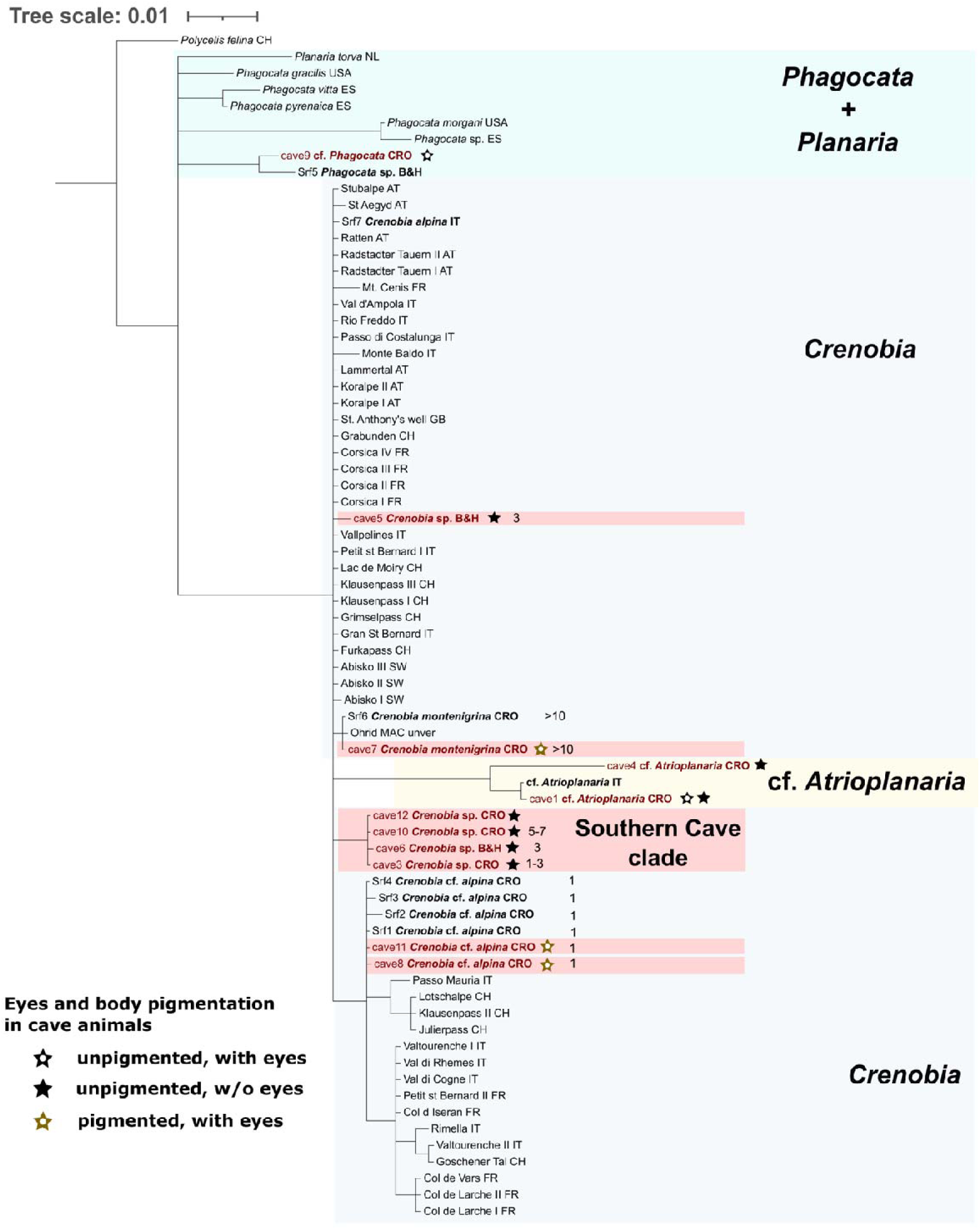

## Appendix 6. ML phylogenetic tree inferred by IQ-TREE from the *COI* gene, using the 1,047 bp long sequence alignment. Bootstrap values lower than 0.95 were collapsed using the ‘delete’ function in iTOL

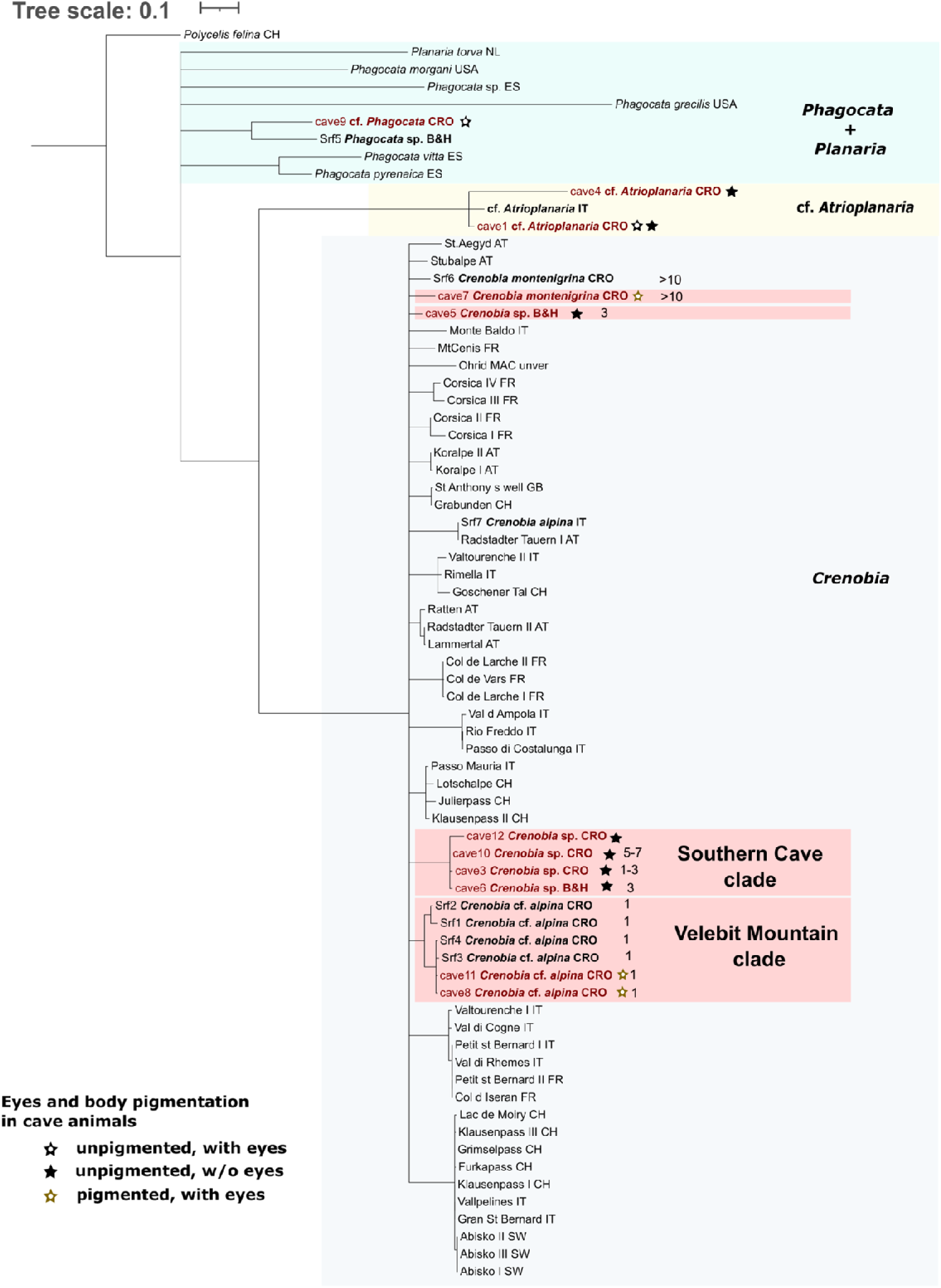

## Appendix 7. BI phylogenetic tree inferred by MrBayes from the *COI* gene, using the 1,047 bp long sequence alignment. Posterior probability values lower than 0.95 were collapsed using the ‘delete’ function in iTOL

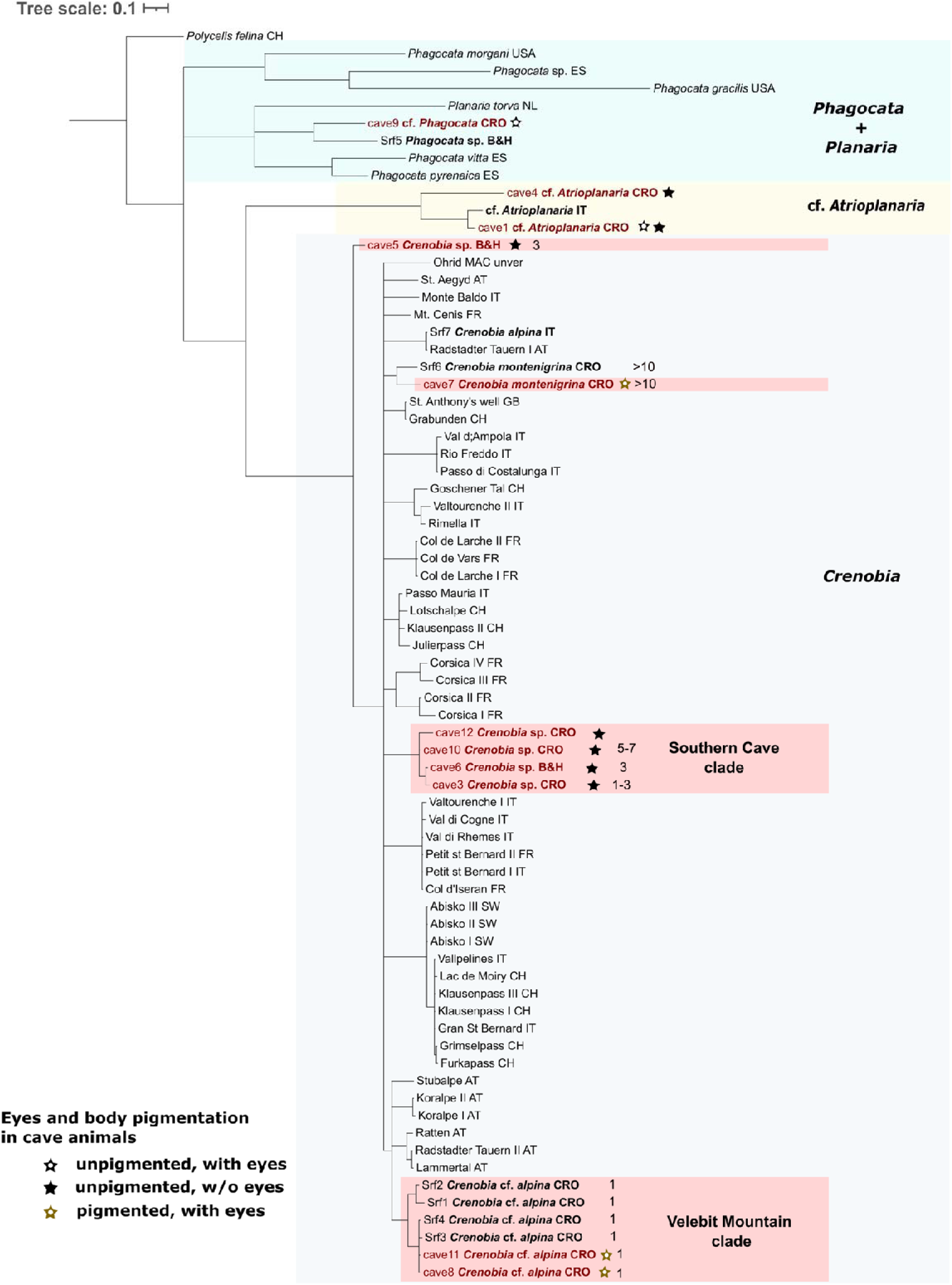

## Appendix 8. Animals from cave1 - Jama na Žutim kokom show variability in the presence of discernible eyes. **A** An animal with both eye spots. **B** An animal with one eye spot. **C** An animal with no discernible eyes

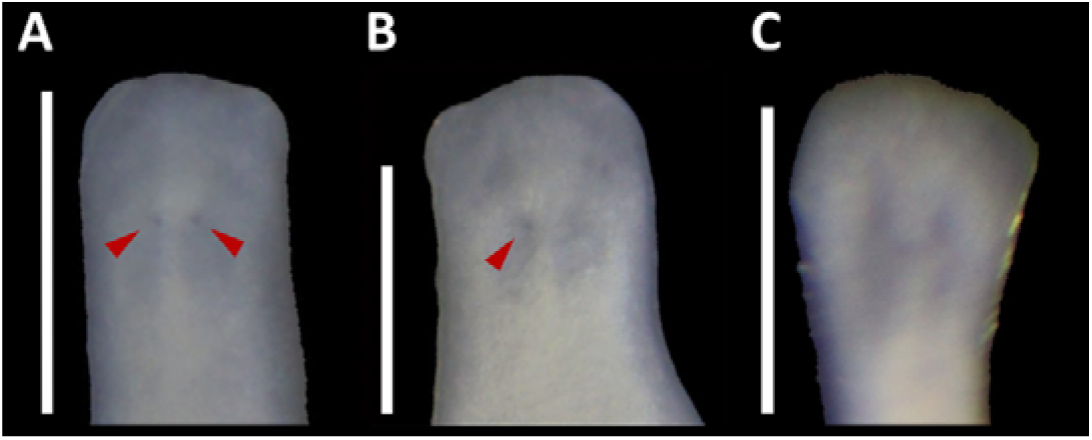

## Appendix 9. BI phylogenetic tree inferred by MrBayes from concatenated *COI + 18S* genes, using ‘full’ alignment dataset, and ‘-allcompat’ function, without pruning of the branches

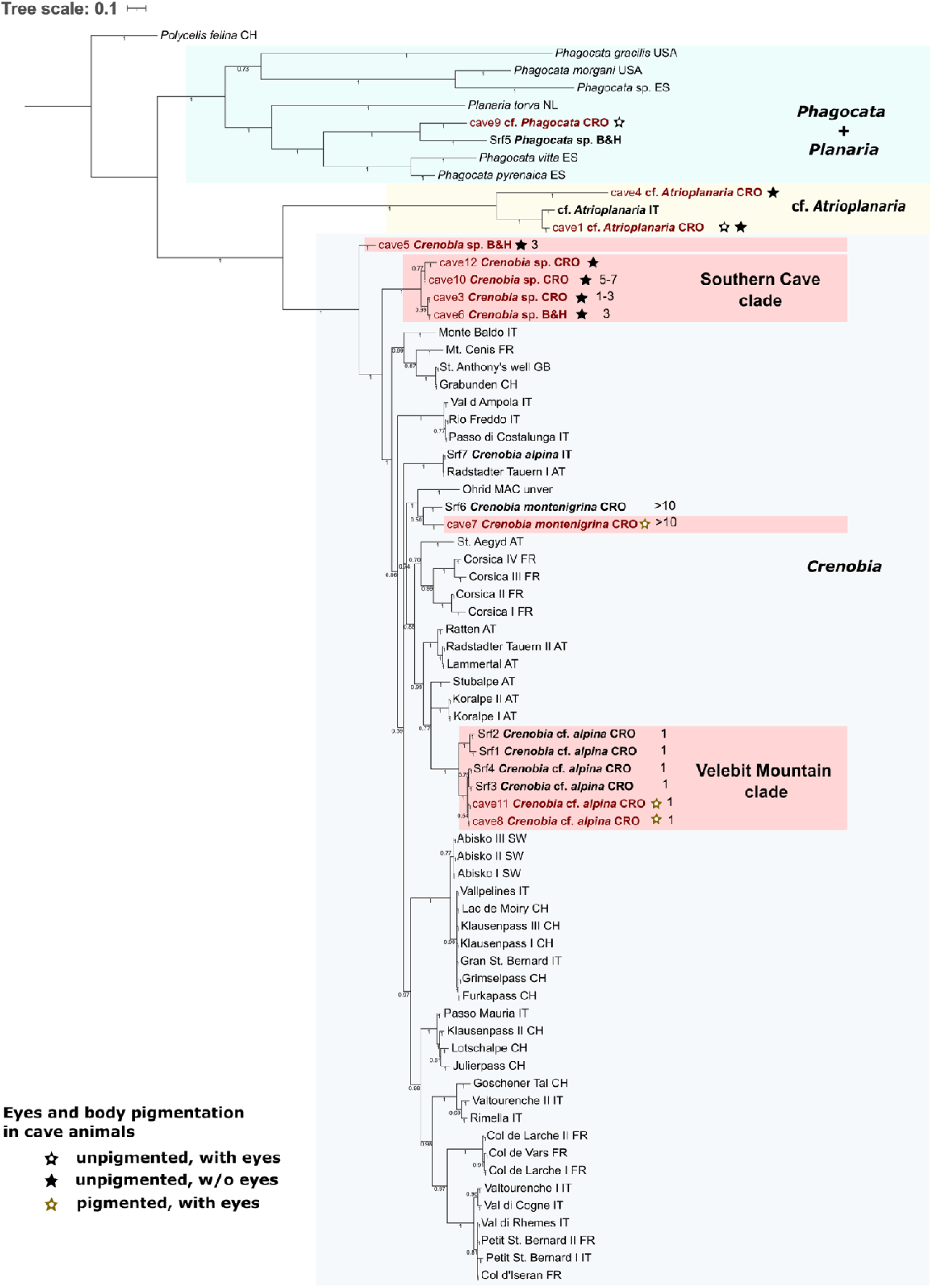

## Notes

### Competing Interest Statement

The authors have declared no competing interest.

